# A dynamic neural network model for predicting risk of Zika in real-time

**DOI:** 10.1101/466581

**Authors:** Mahmood Akhtar, Moritz U.G. Kraemer, Lauren M. Gardner

## Abstract

**Background:** In 2015 the Zika virus spread from Brazil throughout the Americas, posing an unprecedented challenge to the public health community. During the epidemic, international public health officials lacked reliable predictions of the outbreak’s expected geographic scale and prevalence of cases, and were therefore unable to plan and allocate surveillance resources in a timely and effective manner.

**Methods:** In this work we present a dynamic neural network model to predict the geographic spread of outbreaks in real-time. The modeling framework is flexible in three main dimensions i) selection of the chosen risk indicator, *i.e.*, case counts or incidence rate, ii) risk classification scheme, which defines the high risk group based on a relative or absolute threshold, and iii) prediction forecast window (one up to 12 weeks). The proposed model can be applied dynamically throughout the course of an outbreak to identify the regions expected to be at greatest risk in the future.

**Results:** The model is applied to the recent Zika epidemic in the Americas at a weekly temporal resolution and country spatial resolution, using epidemiological data, passenger air travel volumes, vector habitat suitability, socioeconomic and population data for all affected countries and territories in the Americas. The model performance is quantitatively evaluated based on the predictive accuracy of the model. We show that the model can accurately predict the geographic expansion of Zika in the Americas with the overall average accuracy remaining above 85% even for prediction windows of up to 12 weeks.

**Conclusions:** Sensitivity analysis illustrated the model performance to be robust across a range of features. Critically, the model performed consistently well at various stages throughout the course of the outbreak, indicating its potential value at any time during an epidemic. The predictive capability was superior for shorter forecast windows, and geographically isolated locations that are predominantly connected via air travel. The highly flexible nature of the proposed modeling framework enables policy makers to develop and plan vector control programs and case surveillance strategies which can be tailored to a range of objectives and resource constraints.

## Background

The Zika virus, which is primarily transmitted through the bite of infected *Aedes aegypti* mosquitoes (1), was first discovered in Uganda in 1947 (2) from where it spread to Asia in 1960s, where it since has caused small outbreaks. In 2007 ZIKV caused an island wide outbreak in Yap Island, Micronesia (3), followed by outbreaks in French Polynesia (4) and other Pacific islands between 2013–2014 where attack rates where up to 70% (5-7). It reached Latin America between late 2013 and early 2014, but was not detected by public health authorities until May 2015 (8) and since affected 48 countries and territories in the Americas (9-11). Since there is no vaccination or treatment available for Zika infections (12, 13), the control of *Ae. aegypti* mosquito populations remains the most important intervention to contain the spread of the virus (14). In order to optimally allocate resources to suppress vector populations, it is critical to accurately anticipate the occurrence and arrival time of arboviral infections to detect local transmission (15).

Whereas for dengue, the most common arbovirus infection, prediction has attracted wide attention from researchers employing statistical modelling and machine learning methods to guide vector control (16-29), such real-time machine learning based models do not yet exist for Zika virus. Early warning systems for Thailand, Indonesia, Ecuador and Pakistan have been introduced and are currently in use (30-34). In addition to conventional predictions based on epidemiological and meteorological data (20, 35, 36), more recent models have successfully incorporated search engines (37, 38), land use (39), human mobility information (40, 41) and spatial dynamics (42-44), and various combinations of the above (45) to improve predictions. Whereas local spread may be mediated by overland travel, continent wide spread is mostly driven by air passenger travel between climatically synchronous regions (46-52).

The aims of our work are to 1) present recurrent neural networks for time ahead predictive modelling as a highly flexible tool for outbreak prediction, and 2) implement and evaluate the model performance for the Zika epidemic in the Americas. The application of neural networks for epidemic risk forecasting has previously been applied to dengue forecasting and risk classification (53-58), detection of mosquito presence (59), temporal modeling of the oviposition of *Aedes aegypti* mosquito (60), *Aedes* larva identification (61), and epidemiologic time-series modeling through fusion of neural networks, fuzzy systems and genetic algorithms (62). Recently, Jian *et al* (63) performed a comparison of different machine learning models to map the probability of Zika epidemic outbreak using publically available global Zika case data and other known covariates of transmission risk. Their study provides valuable insight into the potential role of machine learning models for understanding Zika transmission; however, it is static in nature, *i.e.*, it does not account for time-series data, and did not account for human mobility, both of which are incorporated in our modelling framework.

Here, we apply a dynamic neural network model for N-week ahead prediction for the 2015-2016 Zika epidemic in the Americas. The model implemented in this work relies on multi-dimensional time-series data at the country (or territory) level, specifically epidemiological data, passenger air travel volumes, vector habitat suitability for the primary spreading vector *Ae. aegypti*, socioeconomic and population data. The modeling framework is flexible in three main dimensions: 1) the preferred risk indictor can be chosen by the policy maker, *e.g*., we consider outbreak size and incidence rate as two primary indicators of risk for a region, 2) five risk classification schemes are defined, where each classification scheme varies in the (relative or absolute) threshold used to determine the set of countries deemed “high risk”, and 3) it can be applied for a range of forecast windows (1 – 12 weeks). Model performance and robustness is evaluated for various combinations of risk indicator, risk classification level, and forecasting windows. Thus, our work represents the first flexible framework of neural networks for epidemic risk forecasting, that allows policy makers to evaluate and weigh the trade-off in prediction accuracy between forecast window and risk classification schemes. Given the availability of the necessary data, the modelling framework proposed here can be applied in real time to future outbreaks of Zika, and other similar vector-borne outbreaks.

## Materials and Methods

### Data

The model relies on socioeconomic, population, epidemiological, travel and mosquito vector suitability data. All data is aggregated to the country level and provided for all countries and territories in the Americas. Each data set and corresponding processing is described in detail below, and summarized in Table 1. All input data is available as Additional files 1-11.

**Table 1.**
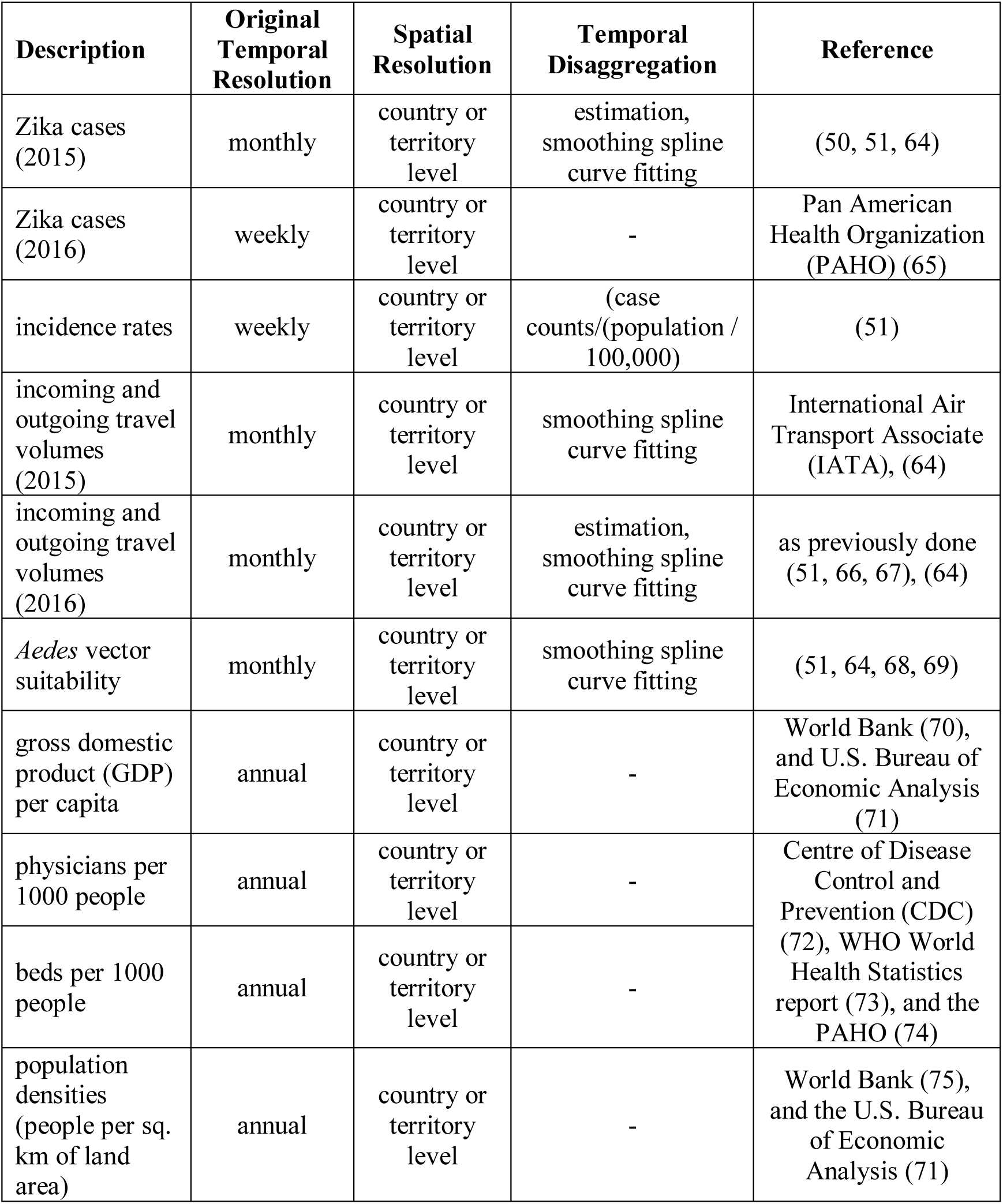
Summary of input data

### EpidemiologicalData

Weekly Zika infected cases for each country and territory in the Americas were extracted from the Pan American Health Organization (PAHO) (65), as described in previous studies (48, 51) (data available: github.com/andersen-lab/Zika-cases-PAHO). The epidemiological weeks 1 - 78 are labeled herein as EPI weeks, corresponding to the dates 29-Jun-2015 - 19-Dec-2016, respectively. Although Zika cases in Brazil were reported as early as May 2015, no case data is available for all of 2015 from PAHO because the Brazil Ministry of Health did not declare the Zika cases and associated neurological and congenital syndrome as notifiable conditions until 17 February of 2016 (65). The missing numbers of cases from July to December 2015 for Brazil were estimated based on the positive correlation between *Ae. aegypti* abundance (described below) and reported case counts as has been done previously (50, 51). We used smoothing spline (64) to estimate weekly case counts from the monthly reported counts. The weekly country level case counts (Figure 1A) were divided by the total population / 100,000, as previously described (51), to compute weekly *incidence rates* (Figure 1B).

**Fig 1.**
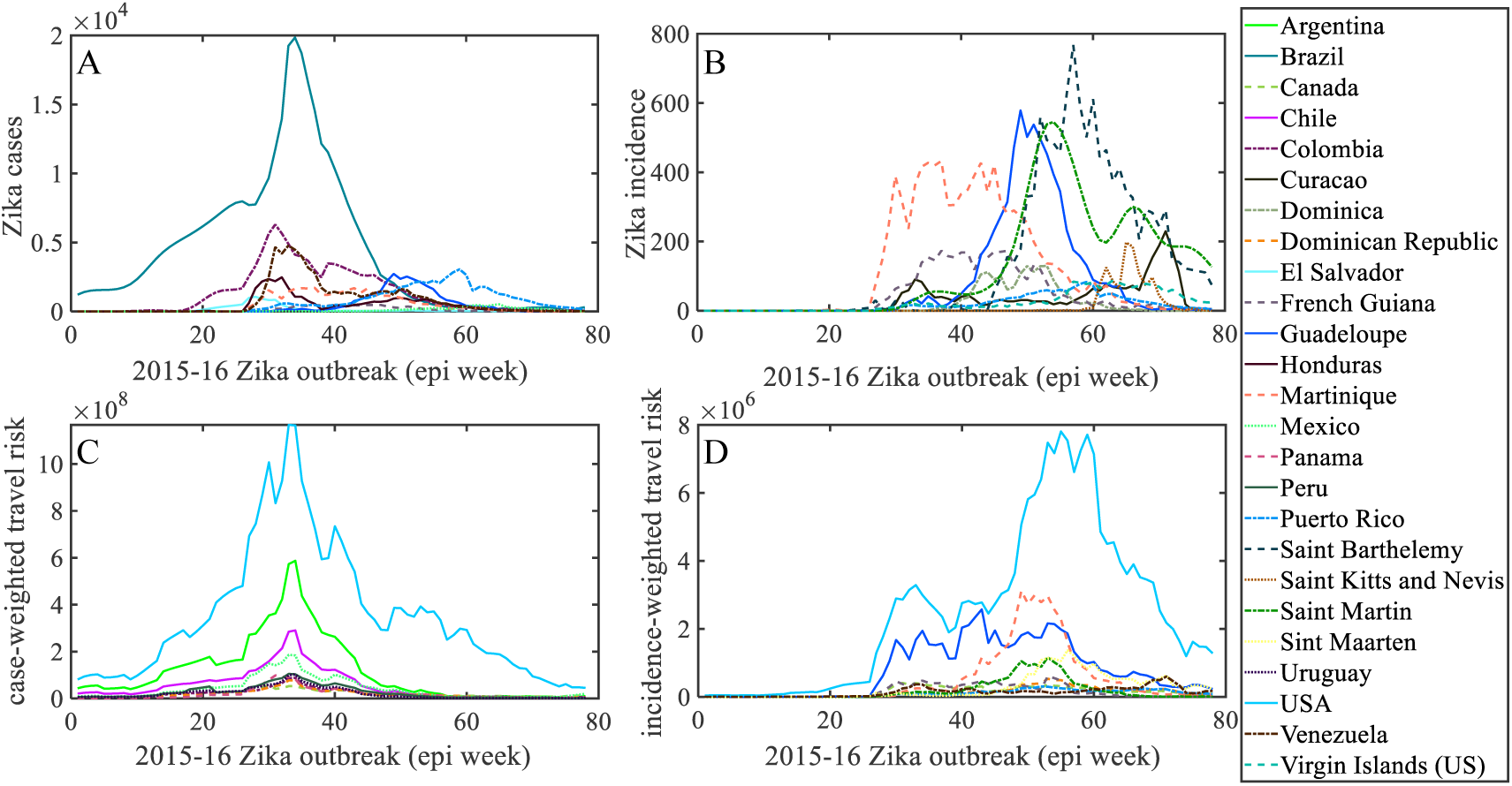
Weekly distribution of case and connectivity-risk variables. (A) Zika cases (B) incidence rates in the Americas, (C) case-weighted travel risk 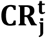, and (D) incidence weighted travel risk 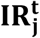, for top 10 ranked countries and territories in the Americas for each respective variable.

### TravelData

Calibrated monthly passenger travel volumes for each airport-to-airport route in the world were provided by the International Air Transport Associate (IATA) (76), as previously used in (51, 66). The data includes origin, destination and stopover airport paths for 84% of global air traffic, and includes over 240 airlines and 3,400 airports. The airport level travel was aggregated to a regional level, to compute monthly movements between all countries and territories in the Americas. The *incoming* and *outgoing travel volumes* for each country and territory, originally available from IATA at a monthly temporal resolution, were curve fitted, again using smoothing spline method (64) to obtain corresponding weekly volumes to match with the temporal resolution of our model. In this study, data and estimates from 2015 were also used for 2016, as was done previously (51, 66, 67).

### MosquitoSuitability Data

The monthly vector suitability data sets were based on habitat suitability for the principal Zika virus species *Ae. aegypti*, previously used in (51), and initially estimated using original high resolution maps (68) and then enriched to account for seasonal variation in the geographical distribution of *Ae. aegypti* by using time-varying covariate such as temperature persistence, relative humidity, and precipitation as well as static covariates such as urban versus rural areas. The monthly data was translated into weekly data using a smoothing spline (64).

### Socioeconomic and Human Population Data

For a country, to prevent or manage an outbreak depends on their ability to implement a successful surveillance and vector control programs (77). Due to a lack of global data to quantify vector control at country level, we utilized alternative economic and health related country indicators which have previously been revealed to be critical risk factors for Zika spread (51). A country’s economic development can be measured by the *gross domestic product (GDP)* per capita at purchasing power parity (PPP), in international dollars. The figures from World Bank (70) and the U.S. Bureau of Economic Analysis (71) were used to collect GDP data for each country. The number of *physicians* and the number of hospital *beds* per 10,000 people were used to indicate the availability of health infrastructure in each country. These figures for U.S. and other regions in the Americas were obtained from the Centre of Disease Control and Prevention (CDC) (72), WHO World Health Statistics report (73), and the PAHO (74). Finally, the human *population densities* (people per sq. km of land area) for each region were collected from World Bank (75) and the U.S. Bureau of Economic Analysis (71).

### Connectivity-risk Variables

In addition to the raw input variables, novel connectivity-risk variables are defined and computed for inclusion in the model. These variables are intended to capture the risk posed by potentially infected travelers arriving at a given destination at a given point in time, and in doing so, explicitly capture the dynamic and heterogeneity of the air-traffic network in combination with real-time outbreak status. Two variables are chosen, hereafter referred to as *case-weighted travel risk* and *incidence-weighted travel risk*, as defined in equations (1.a) and (1.b), respectively.

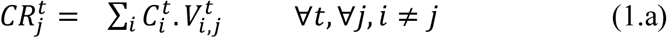

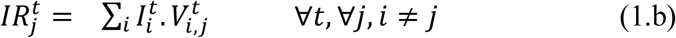

For each region *j* at time *t*, 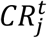 and 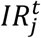 are computed as the sum of product between passenger volume traveling from origin *i* into destination *j* at time 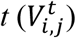 and the state of the outbreak at origin *i* at time t, namely reported cases, 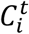, or reported incidence rate, 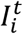. Each of these two variables is computed for all 53 countries or territories for each of the 78 epidemiological weeks. The two dynamic variables, 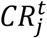 and 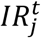, are illustrated in Figure 1C and 1D, below the raw case counts and incidence rates, respectively.

### Neural Network Model

A class of neural architectures based upon Nonlinear Auto Regressive models with eXogenous inputs (NARX) known as NARX neural networks (78-80) is employed herein due to its suitability for modeling of a range of nonlinear systems and computational capabilities equivalent to Turing machines (81). The NARX networks, as compared to other recurrent neural network architectures, require limited feedback (i.e., feedback from the output neuron rather than from hidden states) and converge much faster with a better generalization (81, 82). The NARX model can be formalized as follows (81):

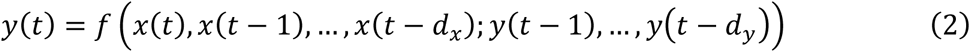

where *x*(*t*) and *y*(*t*) denote, respectively, the input and output (or target that should be predicted) of the model at discrete time *t*, while *d*_*x*_ and *d*_*y*_ (with *d*_*x*_ ≥ 1, *d*_*y*_ ≥ 1, and *d*_*x*_ ≤ *d*_*y*_) are input and output delays called memory orders (Figure 2). In this work, a NARX model is implemented to provide *N-*step ahead prediction of a time series, as defined below:

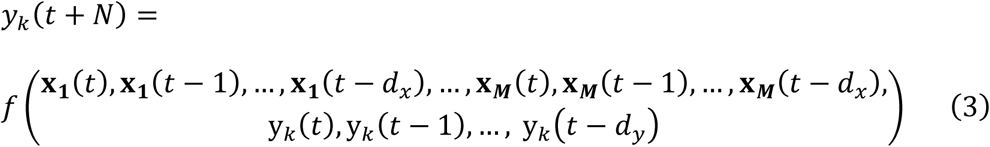

**Fig 2.**
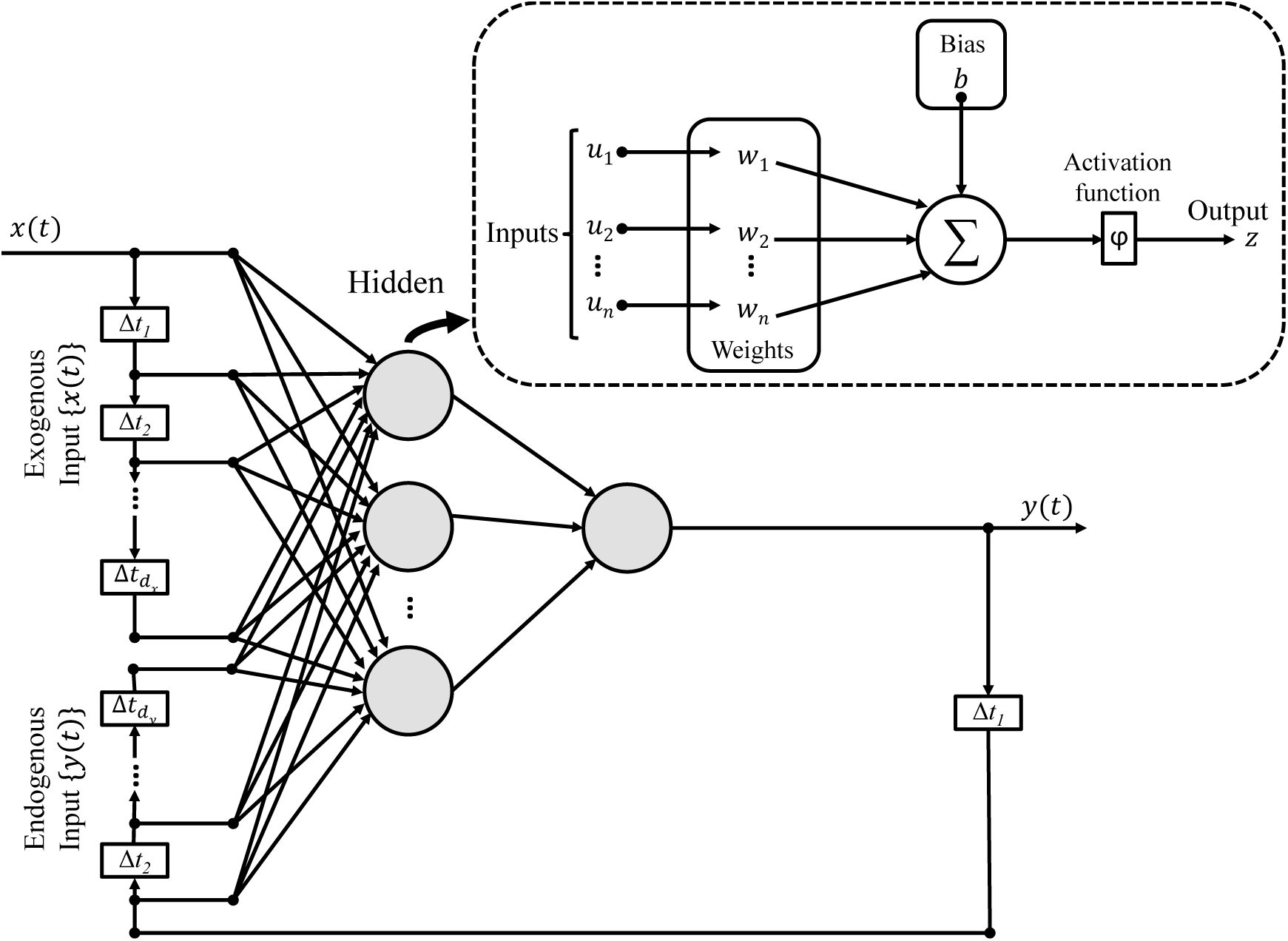
Schematic of NARX network with **d**_**x**_ input and **d**_**y**_ output delays: Each neuron produces a single output based on several real-valued inputs to that neuron by forming a linear combination using its input weights and sometimes passing the output through a nonlinear activation function: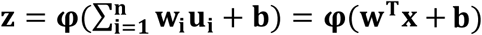, where **w** denotes the vector of weights, **u** is the vector of inputs, **b** is the bias and **φ** is a linear or nonlinear activation function (e.g., Linear, Sigmoid, and Hyperbolic tangent (88)).

Here, *y*_*k*_(*t* + *N*) is the risk classification predicted for the *k*^th^ region *N* weeks ahead (of present time *t*), which is estimated as a function of **x**_***m***_(*t*) inputs from all *m* = 1, 2, …, *M* regions for *d*_*x*_ previous weeks, and the previous risk classification state, *y*_*k*_(*t*) for region *k* for *d*_*y*_ previous weeks. The prediction model is applied at time *t*, to predict for time *t*+*N*, and therefore relies on data available up until week *t.* That is, to predict outbreak risk for epidemiological week X, *N*-weeks ahead, the model is trained and tested using data available up until week (X – N). For example, 12-week ahead prediction for Epi week 40, is performed using data available up to week 28. The function *f*(·) is an unknown nonlinear mapping function that is approximated by a Multilayer Perceptron (MLP) to form the NARX recurrent neural network (79, 80). In this work, series-parallel NARX neural network architecture is implemented in Matlab R2018a (The MathWorks, Inc., Natick, Massachusetts, United States) (83).

In the context of this work, the desired output, *y*_*k*_(*t* + *N*), is a binary risk classifier, *i.e*., classifying a region *k* as high or low risk at time at time *t*+*N*, for each region, *k, N* weeks ahead (of *t*). The vector of input variables for region *m* at time *t* is **x**_***m***_(*t*), and includes both static and dynamic variables. We consider various relative (R) and absolute (A) thresholds to define the set of “high risk” countries at any point in time. We define relative risk thresholds that range uniformly between 10% and 50%, where the 10% scheme classifies the 10% of countries reporting the highest number of cases (or highest incidence rate) during a given week as high risk, and the other 90% as low risk, similar to (45). The relative risk schemes are referred herein as R=0.1, R=0.2, R=0.3, R=0.4, and R=0.5. It is worth noting, for a given percentile, *e.g.*, R=0.1, the relative risk thresholds are dynamic and vary week to week as a function of the scale of the epidemic, while the size of the high risk group remains fixed over time, e.g., 10% of all countries. We also consider absolute thresholds, which rely on case incidence rates to define the “high risk” group. Five absolute thresholds are selected based on the distribution of incidence values over all countries and the entire epidemic. Specifically, the 50th, 60th, 70th, 80th and 90th percentiles were chosen, and are referred herein as A=50, A=60, A=70, A=80, and A=90. These five thresholds correspond to weekly case incidence rates of 0.43, 1.47, 4.05, 9.5 and 32.35 (see Additional file 12: Figure S1), respectively. In contrast to the relative risk scheme, under the absolute risk scheme for a given percentile, *e.g*., A=90, the threshold remains fixed but the size of the high (and low) risk group varies week to week based on the scale of the epidemic. The fluctuation in group size for each threshold is illustrated in Additional file 12: Figure S1 for each classification scheme, A=50 to A=90. Critically, our prediction approach differs from (45), in that our model is trained to predict the risk level directly, rather than predict the number of cases, which are post-processed into risk categories. The performance of the model is evaluated by comparing the estimated risk level (high or low) to the actual risk level for all locations at a specified time. The actual risk level is simply defined at each time period *t* during the outbreak by ranking the regions based on to the number of reported case counts (or incidence rates), and grouping them into high and low risk groups according to the specified threshold and classification scheme.

The static variables used in the model include GDP PPP, population density, number of physicians, and number of hospital beds for each region. The dynamic variables include mosquito vector suitability, outbreak status (both reported case counts and reported incidence rates), total incoming travel volume, total outgoing travel volume, and the two connectivity-risk variables defined as in Equations (1.a) & (1.b), again for each region. Before applying to the NARX model, all data values are normalized to the range [0, 1].

A major contribution of this work is the flexible nature of the model, which allows policy makers to be more or less risk averse in their planning and decision making. Firstly, the risk indicator can be chosen by the modeler; in this work we consider two regional risk indicators, i) the number of reported cases and ii) incidence rate. Second, we consider a range of risk classification schemes, which define the set of high-risk countries based on either a relative or absolute threshold that can be chosen at the discretion of the modeler, *i.e.*, R=0.1, 0.2, 0.3, 0.4, 0.5, and A=90, 80, 70, 60, 50. Third, the forecast window, *N*, is defined to range from *N* = 1, 2, 4, 8 and 12 weeks. Subsequently, any combination of risk indicator, risk classification scheme and forecasting window can be modelled.

In initial settings of the series-parallel NARX neural network, a variety numbers of hidden layer neurons and numbers of tapped delay lines (Eq. (2)) were explored for training and testing of the model. Sensitivity analysis revealed minimal difference in performance of the model under different settings. Therefore, for all experiments presented in this work, the numbers of neural network hidden layer neurons and tapped delay lines are kept constant as two and four, respectively.

To train and test the model, the actual risk classification for each region at each week during the epidemic, *y*_*k*_(*t*), was used. For each model run, e.g., a specified risk indicator, risk classification scheme and forecasting window, the input and target vectors are randomly divided into three sets:

1. 70% for training, to tune model parameters minimizing the mean square error between the outputs and targets,
2. 15% for validation, to measure network generalization and to prevent overfitting, by halting training when generalization stops improving (i.e., mean square error of validation samples starts increasing), and
3. 15% for testing, to provide an independent measure of network performance during and after training.

The performance of the model is measured using two metrics: 1) prediction accuracy (ACC) and 2) receiver operating characteristic (ROC) curves. Prediction accuracy is defined as ACC = (TP + TN) / (TP + FP + TN + FN), where true positive (TP) is the number of high risk locations correctly predicted as high risk, false negative (FN) is the number of high risk locations incorrectly predicted as low risk, true negative (TN) is the number of low risk locations correctly predicted as low risk, and false positive (FP) is the number of low risk locations incorrectly predicted as high risk. The second performance metric, ROC curve (84), explores the effects on TP and FP as the position of an arbitrary decision threshold is varied, which in the context of this prediction problem distinguished low and high risk locations. ROC curve can be characterized as a single number using the area under the ROC curve (AUC), with larger areas having an AUC that approaches one indicating a more accurate detection method. In addition to quantifying model performance using these two metrics, we evaluate the robustness of the predictions by comparing the ACC across multiple runs that vary in their selection of testing and training sets (resulting from the randomized sampling).

## Results

The model outcome reveals the set of locations expected to be at high risk at a specified date in the future, *i.e., N* weeks ahead of when the prediction is made. We apply the model for all epidemiological weeks throughout the epidemic, and evaluate performance under each combination of i) risk indicator, ii) classification scheme, and iii) forecast window. For each model run, both ACC and ROC AUC are computed.

### Model Performance

Figures 3 and 4 exemplify the output of the proposed model. Figure 3 illustrates the model predictions at a country-level for a *4*-week prediction window, specifically for Epi week 40, *i.e*., using data available up until week 36. Figure 3A illustrates the actual risk percentile each country is assigned to in week 40, based on reported case counts. The results presented in the remaining panels of Figure 3 reveal the risk level (high or low) predicted for each country under the five relative risk classification schemes, namely (B) R=0.1, (C) R=0.2, (D) R=0.3, (E) R=0.4, and (F) R=0.5, and whether or not it was correct. For Panels (B)-(E), green indicates a correctly predicted low risk country (TN), light grey indicates an incorrectly predicted high risk country (FP), dark grey indicates an incorrectly predicted low risk country (FN), and the remaining color indicates a correctly predicted high risk country (TP). The inset highlights the results for the *Caribbean* islands. The figure also presents the average ACC over all regions and ACC for just the Caribbean region (grouped similar to (10)) for each classification scheme.

**Fig 3.**
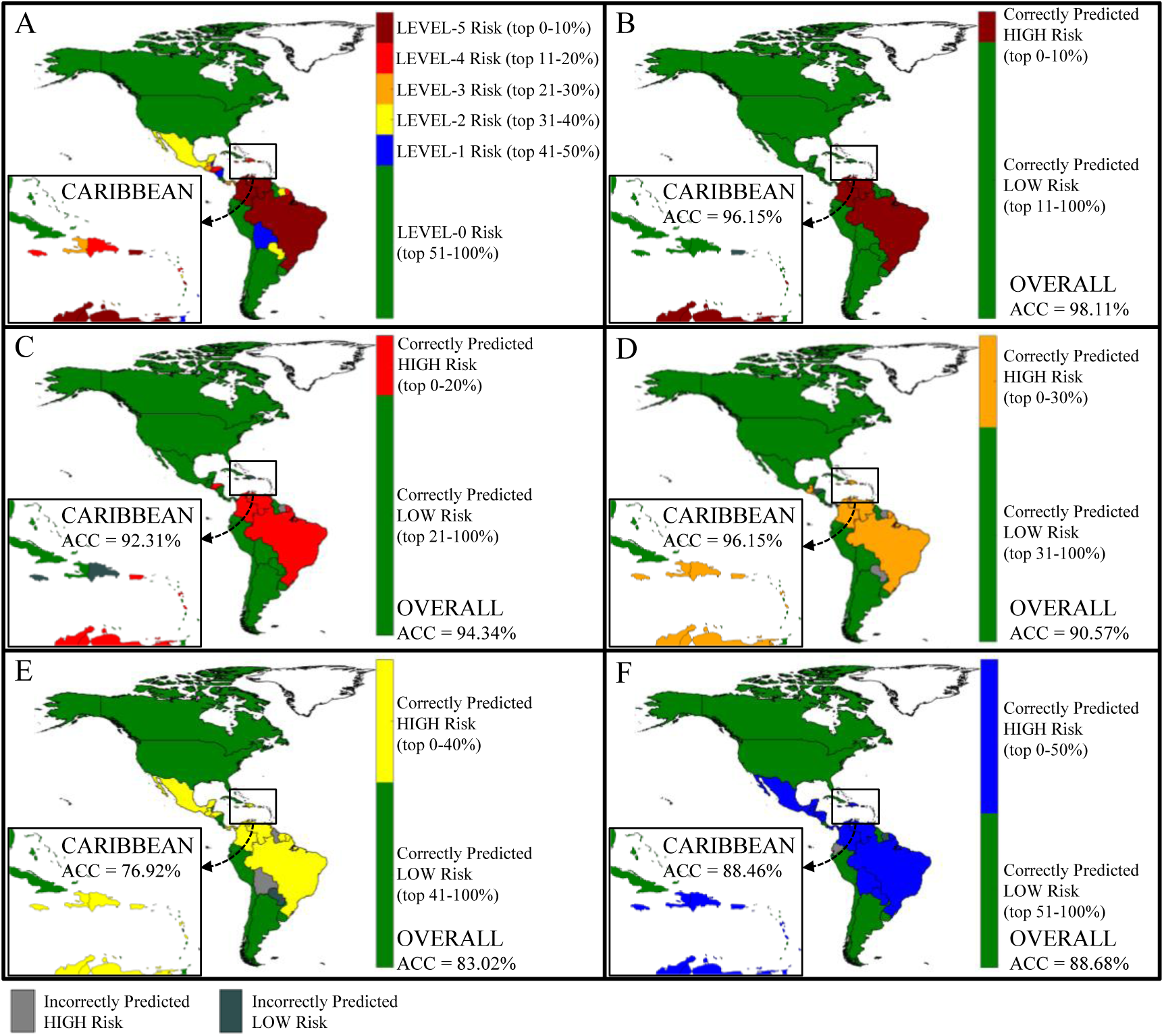
Country prediction accuracy by relative risk level. Panel (A) illustrates the actual relative risk level assigned to each country at Epi week 40 for a fixed forecast window, N=4. Panels (B)-(E) each corresponds to a different classification scheme, specifically (B) R=0.1, (C) R=0.2, (D) R=0.3, (E) R=0.4, and (F) R=0.5. The inset shown by the small rectangle highlights the actual and predicted risk in Caribbean islands. For Panels (B)-(E), green indicates a correctly predicted low risk country, light grey indicates an incorrectly predicted high risk country, and dark grey indicates an incorrectly predicted low risk country. The risk indicator used is case counts.

**Fig 4.**
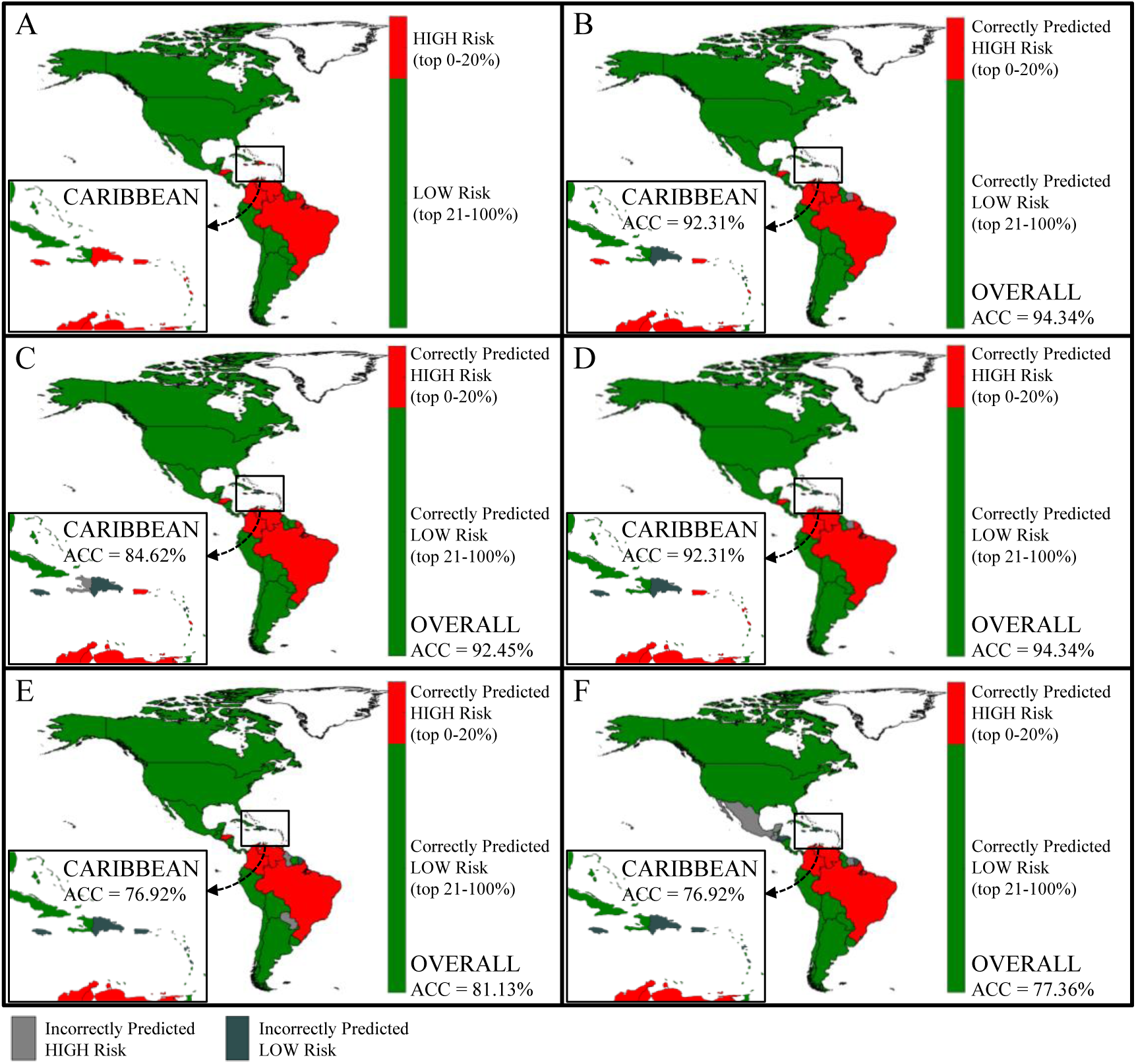
Country prediction accuracy by forecast window. Panel (A) illustrates the actual relative risk level assigned to each country at Epi week 40 for a fixed classification scheme, R=0.2. Panels (B)-(E) each corresponds to different forecast windows, specifically (B) N=1, (C) N=2, (D) N=4, (E) N=8, and (F) N=12. The inset shown by the small rectangle highlights the actual and predicted risk in Caribbean islands. For Panels (B)-(E), the red indicates a correctly predicted high risk country and green indicates a correctly predicted low risk country. Light grey indicates an incorrectly predicted high risk country, and dark grey indicates an incorrectly predicted low risk country. The risk indicator used is case counts.

Figure 4 illustrates the model predictions at a country-level for varying prediction windows, and a fixed classification scheme of R=0.2, again for Epi week 40. Figure 4A illustrates the actual risk classification (high or low) each country is assigned to in Epi week 40, based on reported case counts. The results presented in the remaining panels of Figure 4 reveal the risk level (high or low) predicted for each country under the five forecasting windows, specifically (B) N=1, (C) N=2, (D) N=4, (E) N=8, and (F) N=12, and whether or not it was correct. For Panels (B)-(E), red indicates a correctly predicted high risk country (TP), green indicates a correctly predicted low risk country (TN), light grey indicates an incorrectly predicted high risk country (FP), dark grey indicates an incorrectly predicted low risk country (FN). The inset highlights the results for the *Caribbean* islands. Similar to Figure 3, for each forecast window, the reported ACC is averaged both over all regions and for just the Caribbean.

The model’s performance and sensitivity to the complete range of input parameters is summarized in Additional file 13: Table S2. ACC is presented for each combination of risk indicator (case count and incidence rate), classification scheme (i.e., R = 0.1, 0.2, 0.3, 0.4, 0.5 and A = 90, 80, 70, 60, 50) and forecast window (i.e., *N* = 1, 2, 4, 8 and 12), for selected Epi weeks throughout the epidemic. ROC AUC (averaged over all locations and all EPI weeks) is computed for all combinations of risk indicator (case count and incidence rate), classification scheme (i.e., R = 0.1, 0.2, 0.3, 0.4, 0.5 and A = 90, 80, 70, 60, 50) and forecast window (i.e., *N* = 1, 2, 4, 8 and 12).

Figures 5 and 6 illustrate trends in the model performance as a function of classification scheme and forecast window, aggregated over space and time. Specifically, Figure 5 reveals the model performance (ACC, averaged over all locations and all EPI weeks) for each combination of risk classification scheme (i.e., R = 0.1, 0.2, 0.3, 0.4 and 0.5) and forecast window (i.e., *N* = 1, 2, 4, 8 and 12). The aggregated ROC curves (averaged over all locations and all epidemiological weeks) for R=0.4 are presented in Figure 6, and reveal the (expected) increased accuracy of the model as the forecast window is reduced. The ROC AUC results are consistent with ACC results presented in Figure 5, highlighting the superior performance of the 1 and 2 week ahead prediction capability of the model. The ROC AUC value remains above 0.91 for N=1, 2 and above 0.83 for N=4, both indicating high predictive accuracy of the model. The ROC curves for the other relative risk classification schemes are presented in Additional file 14: Figure S2.

**Fig 5.**
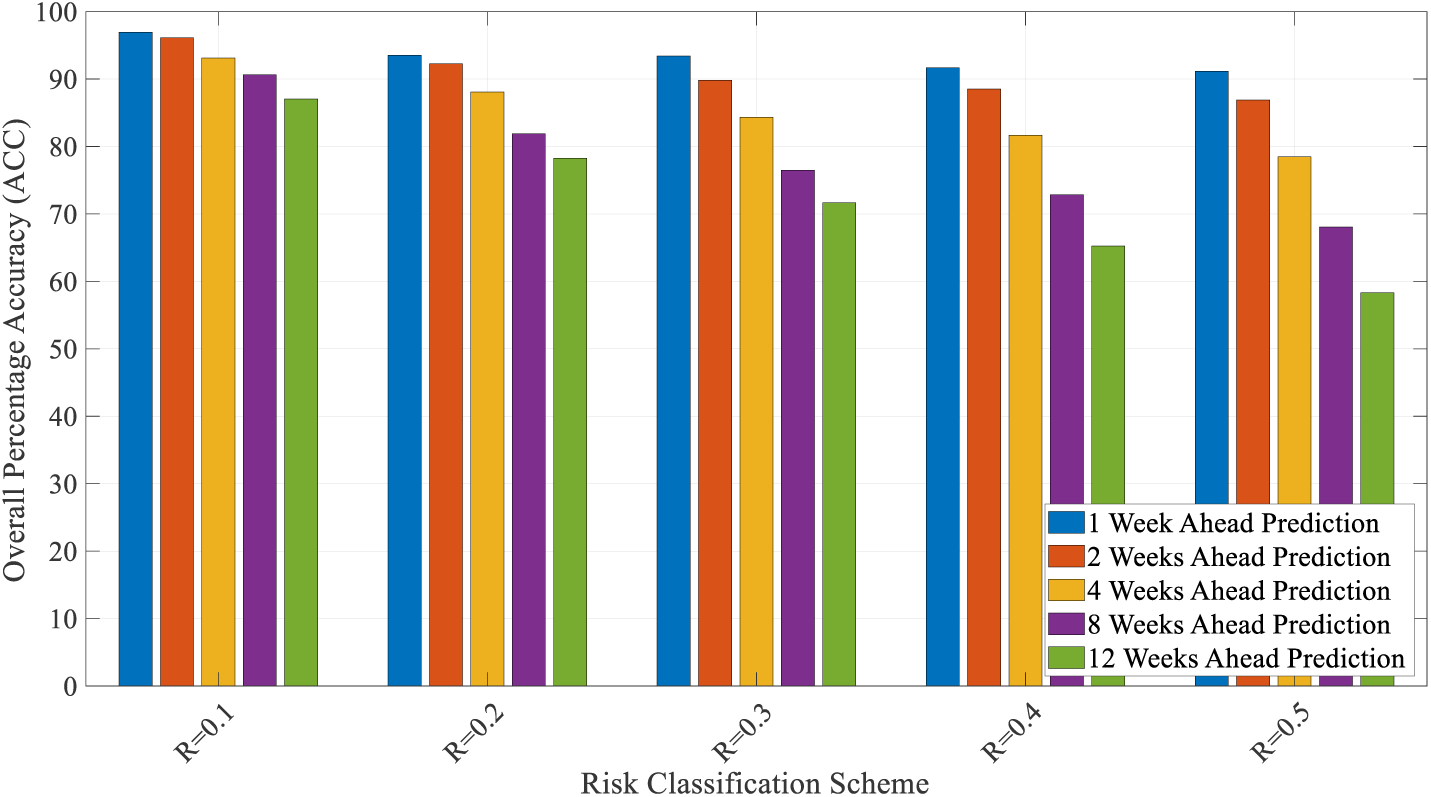
Aggregate model performance measured by ACC. (averaged over all locations and all weeks) for all combinations of relative risk classification schemes (i.e., R = 0.1, 0.2, 0.3, 0.4 and 0.5) and forecast windows (i.e., *N* = 1, 2, 4, 8 and 12), where the risk indicator is case counts.

**Fig 6.**
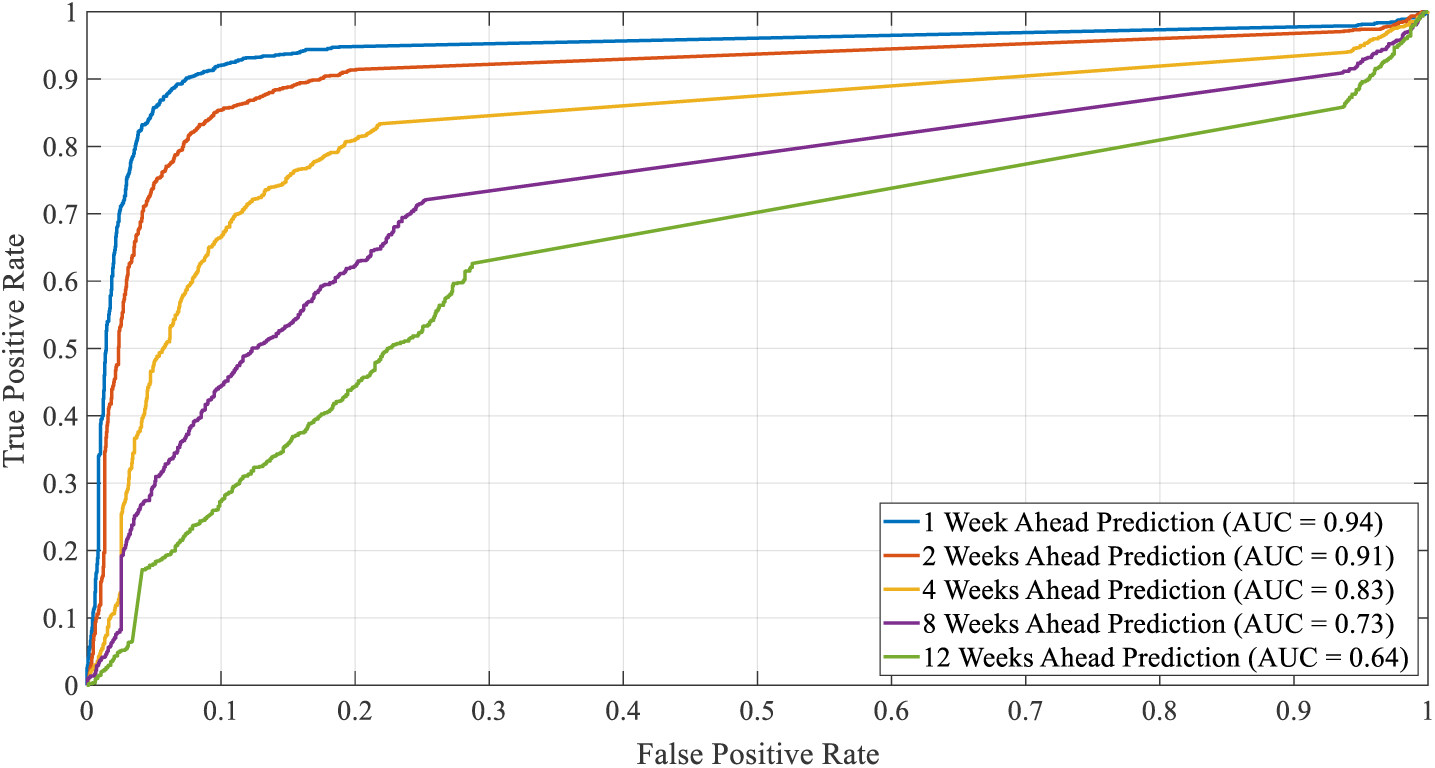
Aggregate model performance measured by ROC AUC. (averaged over all locations and all weeks) for a fixed relative risk classification scheme, *i.e*., R = 0.4, and forecast windows (i.e., *N* = 1, 2, 4, 8 and 12), where the risk indicator is case counts.

### Global and Regional Analysis

We further explore the model’s performance at a regional level by dividing the countries and territories in the Americas into three groups, namely *Caribbean, South America* and *Central America*, as in (10), and compare with the *Global* performance, *i.e.*, all countries. For each group the average performance of the model in terms of ACC was evaluated and presented for each combination of risk indicator (case count and incidence rate), classification scheme (i.e., R = 0.1, 0.2, 0.3, 0.4, 0.5 and A = 90, 80, 70, 60, 50) and forecast window (i.e., *N* = 1, 2, 4, 8 and 12), aggregated over then entire epidemic period (Table 2).

**Table 2.** Summary of Global and Regional Model Performance

| Relative Risk Classification Scheme | Prediction Window Size ($N$ in weeks) | Overall Prediction Accuracy (ACC) | | | | | | | |
| --- | --- | --- | --- | --- | --- | --- | --- | --- | --- |
|  |  | Global |  | Caribbean |  | South America |  | Central America |  |
|  |  | Risk Indicator |  | Risk Indicator |  | Risk Indicator |  | Risk Indicator |  |
|  |  | incidence | cases | incidence | cases | incidence | cases | incidence | cases |
| R=0.1 | 1 | 95.71 | 96.95 | 94.63 | 98.84 | 93.65 | 92.28 | 97.95 | 94.18 |
|  | 2 | 94.29 | 96.12 | 92.90 | 98.86 | 91.68 | 90.78 | 97.01 | 90.67 |
|  | 4 | 91.30 | 93.13 | 89.38 | 97.30 | 87.11 | 84.80 | 95.34 | 84.14 |
|  | 8 | 86.34 | 90.63 | 85.72 | 95.70 | 74.58 | 81.97 | 91.74 | 76.69 |
|  | 12 | 82.57 | 87.05 | 81.75 | 93.59 | 68.63 | 75.94 | 87.99 | 68.14 |
| R=0.2 | 1 | 93.07 | 93.54 | 91.73 | 94.94 | 92.65 | 90.16 | 92.64 | 87.33 |
|  | 2 | 90.01 | 92.27 | 88.30 | 93.93 | 89.37 | 88.60 | 89.26 | 84.68 |
|  | 4 | 84.68 | 88.09 | 82.66 | 89.72 | 82.77 | 84.40 | 82.28 | 76.49 |
|  | 8 | 75.22 | 81.87 | 71.58 | 83.96 | 69.34 | 76.27 | 73.73 | 65.25 |
|  | 12 | 68.96 | 78.25 | 65.01 | 80.92 | 62.75 | 71.30 | 63.73 | 58.09 |
| R=0.3 | 1 | 90.70 | 93.41 | 88.30 | 94.05 | 91.41 | 91.41 | 90.58 | 87.84 |
|  | 2 | 86.74 | 89.82 | 85.27 | 91.06 | 86.68 | 86.68 | 84.15 | 80.46 |
|  | 4 | 80.85 | 84.31 | 77.10 | 85.36 | 82.63 | 79.38 | 78.73 | 72.76 |
|  | 8 | 70.10 | 76.46 | 64.73 | 77.31 | 69.34 | 71.34 | 66.31 | 58.05 |
|  | 12 | 63.37 | 71.66 | 56.86 | 70.66 | 62.39 | 64.88 | 57.35 | 56.86 |
| R=0.4 | 1 | 90.46 | 91.68 | 88.25 | 91.31 | 93.03 | 90.54 | 87.84 | 86.64 |
|  | 2 | 86.79 | 88.52 | 84.62 | 88.24 | 89.76 | 86.30 | 83.10 | 81.51 |
|  | 4 | 79.36 | 81.67 | 76.41 | 81.97 | 83.04 | 75.44 | 72.57 | 71.83 |
|  | 8 | 68.47 | 72.85 | 61.67 | 71.12 | 73.19 | 71.19 | 62.29 | 54.66 |
|  | 12 | 59.82 | 65.22 | 51.89 | 60.33 | 62.21 | 60.96 | 53.19 | 53.43 |
| R=0.5 | 1 | 89.51 | 91.16 | 89.67 | 90.25 | 87.42 | 88.42 | 85.10 | 90.41 |
|  | 2 | 86.21 | 86.90 | 84.83 | 86.13 | 85.15 | 82.84 | 83.10 | 84.15 |
|  | 4 | 77.67 | 78.46 | 76.29 | 77.55 | 75.44 | 75.71 | 70.71 | 67.72 |
|  | 8 | 66.42 | 68.05 | 61.99 | 65.78 | 69.80 | 68.26 | 53.60 | 47.46 |
|  | 12 | 56.16 | 58.31 | 48.04 | 51.81 | 62.21 | 54.90 | 43.38 | 46.81 |

### Model Robustness

Figure 7A and 7B show how the ACC varies over 10 independent runs of the model. This sensitivity analysis was conducted for all combinations risk indicator, relative risk classification schemes, and selected epidemiological weeks (i.e., week number / starting date: 30 / 18-Jan-2016, 40 / 28-Mar-2016, 50 / 6-Jun-2016, 60 / 15-Aug-2016, and 70 / 24-Oct-2016). This time period represents a highly complex period of the outbreak with country level rankings fluctuating substantially, as evidenced in Figure 1. Due to computation time, the sensitivity analysis was evaluated for only the 4-week forecast window. The size of the error bars illustrates the robustness of the proposed modeling framework.

**Fig 7.**
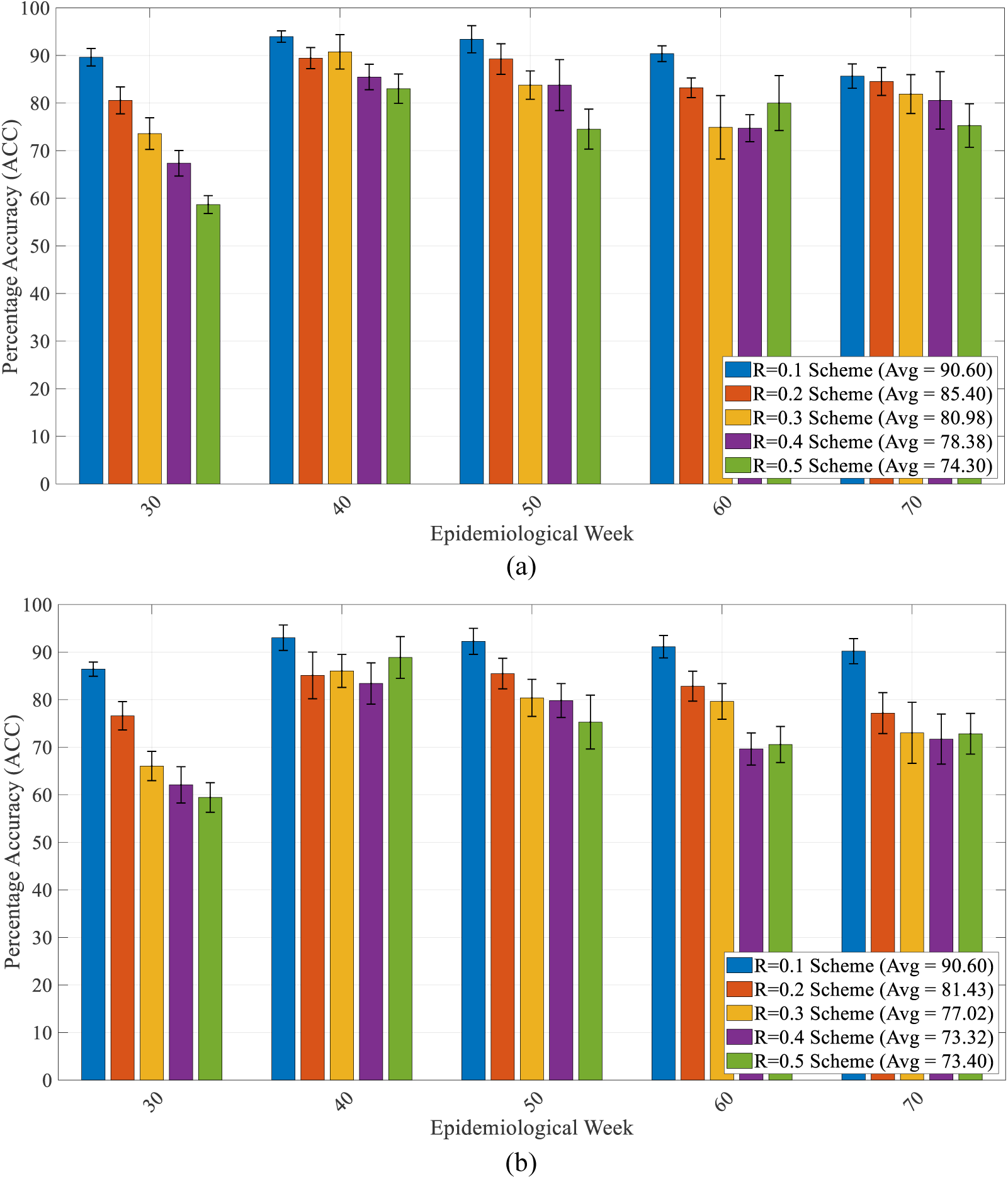
Model performance and robustness. ACC is averaged over all locations for selected epidemiological weeks when risk indicator is (a) case counts and (b) incidence rate, and a fixed forecast windows (i.e., N = 4). The error bars represent the variability in expected ACC across ten runs for each combination.

### NARX Feature Selection

While the NARX framework does not provide assigned weights for each input feature as output, sensitivity analysis can be conducted to help identify the key predictive features. We tested the performance of the NARX framework under three different combinations of input features, with the particular objective of quantifying the role of travel data in our outbreak prediction model. We considered i) a simple ‘baseline’ model using only case count and incidence data, ii) an expanded baseline model that includes case and incidence data, and all non-travel related variables, and iii) the proposed model which includes all features listed in Table 1. The results comparing the performance of these three models with the detailed list of input features for each is provided in Additional file 15: Table S1. The results reveal the case-related data (regional case counts and incidence rates) to be the dominant explanatory variables for predicting outbreak risk in a region, as would be expected. The inclusion of non-travel related variables (regional suitability, regional GDP, regional physicians, regional hospital beds, regional population density) is not shown to improve predictive capability over the baseline model, and in fact, sometime performs worse than the baseline model. In contrast, the inclusion of travel data (weekly case-weighted travel risk, weekly incidence-weighted travel risk, weekly incoming travel volume, weekly outgoing travel volume) is revealed to improve the predictive capability, especially for the shorter prediction windows, with a higher AUC ROC for a majority (20 of the 25) of the scenarios tested. These results support the inclusion of the dynamic travel-related variables, which substantially increase the complexity of the model (inputs), and thus, justifies the use of the NARX framework selected.

## Discussion

Overall, the proposed model is shown to be accurate and robust, especially for shorter prediction windows and higher risk thresholds. As would be expected, the performance of the proposed model decreases as the prediction window increases because of the inherent uncertainty in outbreak evolution over long periods of time. Specifically, the model is almost 80% accurate for 4-week ahead prediction for all classification schemes, and almost 90% accurate for all 2-week ahead prediction scenarios, *i.e.*, the correct risk category of 9 out of 10 locations can always be predicted, indicating strong performance. Although, when the objective is to identify the top 10% of at-risk regions, the average accuracy of the model remains above 87% for prediction up to *12*-weeks in advance. Generally, the model performance is shown to decrease as the risk threshold is reduced, *e.g.*, the size of the high risk group is increased, representing a more risk averse policy. The decrease in performance is likely due to the increased size and fluctuation of the high risk country set over time for lower thresholds. For example, for the absolute risk threshold of A=50, the number of countries classified as high risk fluctuates between 1 and 34 throughout the course of the epidemic, compared with A=90, where the set only ranges from 0 to 12 (see Additional file 12: Figure S1). These results reveal the trade-off between desired forecast window and precision of the high risk group. The quantifiable trade-off between the two model inputs (classification scheme and forecast window) can be useful for policies which may vary in desired planning objectives.

The results in Figures 3 and 4, as well as Table 2 reveal a similar trend at the regional level as was seen at the global level, with a decrease in predictive accuracy as the forecast window increases in length, and the and high risk group increases in size. As shown in Figure 3, the ACC remains above 90% for R < 0.3, indicating superior model performance. For example, at Epi week 40, R = 0.3 and N=4 (using outbreak data and other model variables up to Epi week 36), there were 16 total regions classified as high risk, of which the model correctly identified 13. Furthermore, of the 16 high risk regions, 8 were in the *Caribbean* (i.e., Aruba, Curacao, Dominican Republic, Guadeloupe, Haiti, Jamaica, Martinique, and Puerto Rico), of which the model correctly identified 7. Aruba in the only Caribbean, and Honduras and Panama were the only regions incorrectly predicted as low risk in this scenario; accurately classifying low risk regions is also important (and assuring the model is not too risk averse). For the same scenario, *i.e.*, Epi week 40, R = 0.3 and N=4, all 18 low risk *Caribbean* locations and 17 of the 19 low risk non-*Caribbean* locations were accurately classified by the model. Paraguay and Suriname were the only regions incorrectly predicted as high risk. These results are consistent with the high reported accuracy of the model, i.e., Overall ACC = 90.15%; *Caribbean* ACC = 96.15%. Figure 4 reveals that the performance of model, expectedly, deteriorates as the forecast window increases; however, the average accuracy remains above 80% for prediction up to *8*-weeks ahead, and well about 90% for up to 4-weeks ahead. The prediction accuracy for the Caribbean slightly lags the average performance in the Americas. Specifically, for R=0.2, 5 of the 11 *Caribbean* regions were designated as HIGH risk locations at Epi week 40, *i.e.*, Dominican Republic, Guadeloupe, Jamaica, Martinique, Puerto Rico. For a one-week prediction window, N=1, the model was able to correctly predict 3 of the high risk regions (*i.e.*, Jamaica, Martinique, Puerto Rico), for N=2 it correctly identified two (*i.e.*, Martinique, Puerto Rico), and for N=4, it again correctly identified three (*i.e.*, Guadeloupe, Martinique, Puerto Rico).

However, the model did not correctly predict any high risk locations in the Caribbean at N=8 and N=12 window lengths. This error is due to the low and sporadic reporting of Zika cases in the region around week 30, and the high variability of the outbreak over the 8 and 12 week period. Similar prediction capability is illustrated for R=0.5 (not shown in the figure), in which case out of the 13 *Caribbean* HIGH risk locations, the model correctly identifies all locations at N=1, 2 and 4, 10 of the 13 locations at N=8, and only 1 of the 13 at N=12.

When comparing performance across regions (see Table 2) results reveal the predictive accuracy is best for the *Caribbean* region, while predictions for *Central America* were consistently the worst; the discrepancy in performance between these groups increases as the forecast window increases. The difference in performance across regions can be attributed to the high spatial heterogeneity of the outbreak patterns, the relative ability of air travel to accurately capture connectivity between locations, and errors in case reporting that may vary by region. For example, the *Caribbean*, which consists of more than twice as many locations as any other group, first reported cases around week 25, and remained affected throughout the epidemic. In contrast, *Central America* experienced a slow start to the outbreak (at least according to case reports) with two exceptions, namely Honduras and El Salvador. The large number of affected region in the Caribbean, with more reported cases distributed over a longer time period contributed to the training of the model, thus improving the predictive capability for these regions. Additionally, the geographically isolated nature of Caribbean islands enables air travel to more accurately capture incoming travel risk, unlike countries in Central and South America, where individuals can also move about using alternative modes, which are not accounted for in this study. These factors combined explain the higher predictive accuracy of the model for the Caribbean region, and importantly, helps to identify the critical features and types of settings under which this model is expected to perform best.

Finally, the robustness of the model predictions is illustrated by the short error bars in Figure 7. The model is also demonstrated to perform consistently throughout the course of the epidemic, with the exception of week 30, at which time there was limited information available to train the model, *e.g.*, the outbreak was not yet reported in a majority of the affected countries. Comparing Figure 7A and 7B reveals relatively similar performance for both risk indicators, and Additional File 13: Table 2 demonstrating the model’s flexibility and adaptability with respect to both the risk scheme chosen, *i.e.*, relative or absolute, and the metric used to classify outbreak risk, *i.e.*, number of cases or incidence rate in a region.

### Limitations

There are several limitations of this work. The underlying data on case reporting vary by country and may not represent the true transmission patterns (85). However, the framework presented was flexible enough to account for these biases and we anticipate will only be improved as data become more robust. Additionally, 2015 travel data was used in place of 2016 data, as has been done previously (51, 66, 67), which may not be fully representative of travel behaviour. Furthermore, air travel is the only mode of travel accounted for, thus, additional person movements between country pairs that share land borders are unaccounted for, and as a result, the model likely underestimates the risk posed to some regions. This limitation may partially explain the increased model performance for the geographically isolated Caribbean Islands, which represent a large proportion of ZIKV affected regions. This study does not account for species of mosquitos other than *Ae. Aegypti*, such as *Ae. Albopictus*, which can also spread ZIKV; however, *Ae. Aegypti* are known to be the primary spreading vector, and responsible for the majority of the ZIKV epidemic in the Americas (86). Additionally, alternative non-vector-borne mechanisms of transmission are ignored. Lastly, due to the lack of spatial resolution of case reports, we were limited to make country to country spread estimates. We do however appreciate that there is considerable spatial variation within countries (i.e., northern vs. southern Brazil) and that this may influence the weekly covariates used in this study. We again hypothesise that models will become better as the spatial resolution of available data increases.

## Conclusions

We have introduced a flexible, predictive modelling framework to forecast outbreak risk in real-time that can be scaled and readily applied in future outbreaks. An application of the model was applied to the Zika epidemic in the Americas at a weekly temporal resolution, and country-level spatial resolution, using a combination of population, socioeconomic, epidemiological, travel patterns and vector suitability data. The model performance was evaluated for various risk classification schemes, forecast windows and risk indicators, and illustrated to be accurate and robust across a broad range of these features. First, the model is more accurate for shorter prediction windows and restrictive risk classification schemes. Secondly, regional analysis reveals superior predictive accuracy for the Caribbean, suggesting the model to be best suited to geographically isolated locations that are predominantly connected via air travel. Predicting the spread to areas that are relatively isolated has previously been shown to be difficult due to the stochastic nature of infectious disease spread (87). Thirdly, the model performed consistently well at various stages throughout the course of the outbreak, indicating its potential value at the early stages of an epidemic. The outcomes from the model can be used to better guide outbreak resource allocation decisions, and can be easily adapted to model other vector-borne epidemics.

## Additional files

**Additional file 1: Data (cases).** Country or territory level weekly Zika cases.

**Additional file 2: Data (incidence).** Country or territory level weekly Zika incidence rates.

**Additional file 3: Data (incoming_travel).** Country or territory level, weekly incoming travel volume.

**Additional file 4: Data (outgoing_travel).** Country or territory level weekly outgoing travel volume.

**Additional file 5: Data (suitability).** Country or territory level weekly *Aedes* vector suitability.

**Additional file 6: Data (gdp).** Country or territory level GDP per capita.

**Additional file 7: Data (physicians).** Country or territory level physicians per 1000 people.

**Additional file 8: Data (beds).** Country or territory level beds per 1000 people.

**Additional file 9: Data (pop_density).** Country or territory level population densities (people per sq. km of land area).

**Additional file 10: Data (case_weighted_travel_risk).** Country or territory level weekly case-weighted travel risk.

**Additional file 11: Data (incidence_weighted_travel_risk).** Country or territory level weekly incidence-weighted travel risk.

**Additional file 12: Figure S1. Number of high risk countries each week under all absolute risk classification schemes.** The number of countries classified as high risk each week for each absolute case incidence threshold, ranging from A=50 to A=90. In parentheses is the weekly incidence rate defining the high risk threshold based on the percentile (A) specified.

**Additional file 13: Table S2. Summary of model performance.** ACC is presented for each combination of risk indicator (case count and incidence rate), classification scheme (i.e., R = 0.1, 0.2, 0.3, 0.4, 0.5 and A = 90, 80, 70, 60, 50) and forecast window (i.e., *N* = 1, 2, 4, 8 and 12), for selected Epi weeks throughout the epidemic. ROC AUC (averaged over all locations and all EPI weeks) is computed for all combinations of risk indicator (case count and incidence rate), classification scheme (i.e., R = 0.1, 0.2, 0.3, 0.4, 0.5 and A = 90, 80, 70, 60, 50) and forecast window (i.e., *N* = 1, 2, 4, 8 and 12).

**Additional files 14: Figure S2. Aggregate model performance measured by ROC AUC.** The ROC AUC is averaged over all locations and all weeks, for each relative risk classification scheme, *i.e*., R = 0.1, 0.2, 0.3, 0.4, 0.5 and forecast window *i.e., N* = 1, 2, 4, 8 and 12. For the results shown the risk indicator is case counts.

**Additional file 15: Table S1. Summary of model sensitivity to feature selection.** The ACC and ROC AUC performance of the model is computed and presented under different combinations of input data features. The proposed model is compared against two baseline models; one includes only case (and incidence) data, and the second includes case and all non-travel related data, while the final proposed model includes all features. The results presented are for the absolute risk classification scheme, where the risk indicator is incidence rate.

## Abbreviations

ACC: Prediction accuracy
AUC: Area under the curve
CDC: Centre of disease control and prevention
FN: False negative
FP: False positive
GDP: Gross domestic product
IATA: International air transport associate
MLP: Multilayer perceptron
NARX: Nonlinear autoregressive models with exogenous inputs
PAHO: Pan American health organization
PPP: Purchasing power parity
ROC: Receiver operating characteristic
TN: True negative
TP: True positive
ZIKV: Zika virus

## Ethics approval and consent to participate

Not applicable.

## Consent for publication

Not applicable.

## Availability of data and material

All data used in this study is provided as Additional files.

## Authors’ Contributions

LG and MA conceived the study, designed the experiments, analyzed the model results, and drafted the original manuscript. MA developed the model and performed the computational analysis. MUGK contributed vector distribution data. All authors contributed to data curation and editing of the manuscript. LG supervised the study.

## Acknowledgements

We thank Raja Jurdak and Dean Paini for their inputs and discussion on the model.

